# The impact of selected abiotic factors on zooplankton hatching process through real-time, in-situ observation

**DOI:** 10.1101/2023.01.20.524934

**Authors:** Preyojon Dey, Terence M. Bradley, Alicia Boymelgreen

## Abstract

Current studies on abiotic impacts on marine microorganisms often focus on endpoint analysis (e.g., hatching rates, survival). Here, we demonstrate that a mechanistic understanding can be obtained through real-time measurement of respiration and morphology in controlled microenvironments over extended time periods. As a demonstration, temperature and salinity are chosen to represent critical abiotic parameters that are also threatened by climate change and a target species of *Artemia*, a prominent zooplankton whose reproduction can affect the marine food pyramid. Different temperatures (20, 35, and 30ºC) and salinities (0, 25, 50, and 75 ppt) are shown to significantly alter the duration of hatching stages, metabolic rates, and hatchability. Higher temperatures and moderate salinity boosted metabolic reactivation of latent cysts, while higher temperatures alone sped up the process. Hatchability is inversely related to the duration of the differentiation stage of hatching, which persisted longer at lower temperatures and salinities. Initial oxygen availability affects respiration but not hatchability owing to temperature and salinity interactions.

**Graphical abstract:** 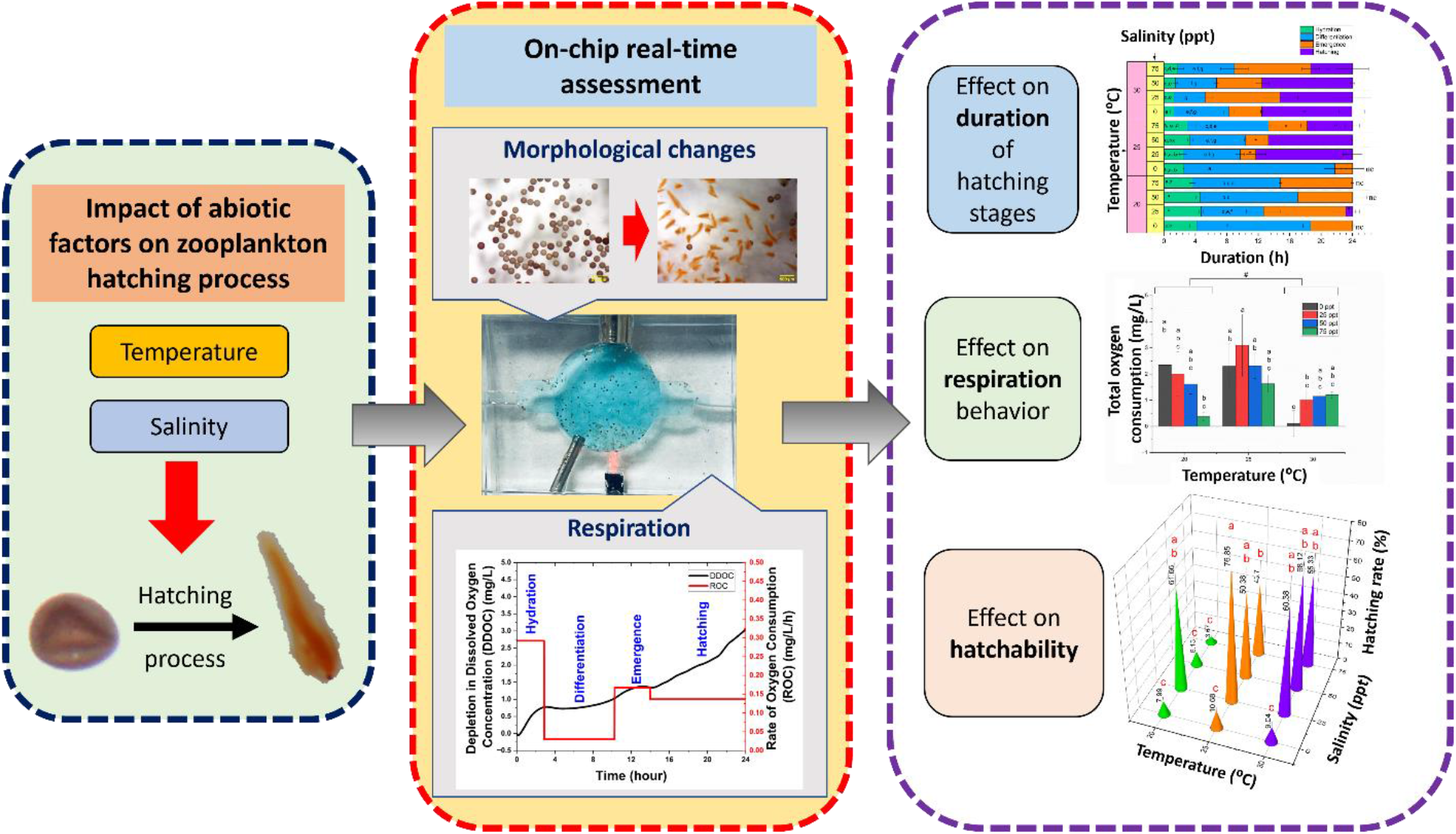

## 1. Introduction

The rising concentration of greenhouse gases caused by anthropogenic activity is modifying the global climate, and one of the immediate results is higher ambient temperature (Hegerl and Cubasch, 1996; Hoegh-Guldberg and Bruno, 2010; Shen et al., 2020). The oceans around the world, acting as heat sinks, absorb the bulk of this heat (Lyman et al., 2010; Nitzbon et al., 2022). Thermal expansion of the oceans as a result of increased heat content, as well as glacier melting, produces a rise in ocean volume, which has a direct effect on ocean water salinity (Cazenave and Llovel, 2010; Wadhams and Munk, 2004) that is further exacerbated by changes in the global water cycle (Douville et al., 2021; Haddeland et al., 2014). The effects of variations in these parameters on reproduction and health – which will have effects not on only individual elements but the overall population (Hiscock et al., 2004; Jaspers et al., 2011; Lalli and Parsons, 1997; Lin et al., 2001; Lushchak, 2011; Orton, 1920) and thus by extension the marine ecosystem as a whole - have been widely studied on organisms ranging from mysid shrimp (*Mysidopsis bahia*) to finfish (leopard grouper; *Mycteroperca rosacea*) (Gracia-López et al., 2004; MCKENNEY JR, 1996). However, most studies are restricted to endpoint analysis (survival rates) or global measurements such as swimming performance, and the precise mechanisms by which the environment affects these biological processes is not well defined.

Here, we aim to obtain a mechanistic understanding of the impacts of changes in the abiotic environment on reproduction through in situ monitoring of the hatching process. *Artemia* – commonly referred to as brine shrimp - has been selected as a model marine species in this study, since it plays an essential role both in the natural marine environment and commercial aquaculture. Specifically, as a major zooplankton, *Artemia* have both direct and indirect effects on fish populations and recruitment (Dettmers et al., 2003; Lomartire et al., 2021; Pershing et al., 2005; van Deurs et al., 2009), serve as an essential source of nitrogen and phosphorous for phytoplankton (Lehman and Sandgren, 1985; Sterner, 2009) and also contribute to the efficiency of the ocean as a biological carbon pump (BCP) by limiting particle export through diel vertical migration (Archibald et al., 2019; Lomartire et al., 2021). Commercially, the high nutritional density and ease of culture makes the species optimal for facilitating feeding of marine larvae with small mouth gape (Camargo et al., 2005; Rocha et al., 2017).

*Artemia* has two alternative reproduction modes. When the environment is favorable (sufficient dissolved oxygen and moderate temperature and salinity), the female *Artemia* produces thin-shelled eggs that complete development and hatch in the uterus (ovoviviparity). In the alternative mode (oviparity), females produce a diapause cyst, with no measurable metabolic activity (endogenous diapause) (Parra and Yúfera, 2001; Wang et al., 2019). In this dormant stage, cysts may be dehydrated through air drying or osmotic water removal, at which point the cysts are quiescent and can survive up to 28 years (Clegg, 1962). The metabolism of the cysts can be resumed -and hatching initiated - when the environmental conditions are favorable (Liu et al., 2009), where the primary abiotic environmental parameters affecting hatchability are water temperature and salinity (Arun et al., 2017; Browne and Wanigasekera, 2000; Lenz and Browne, 2018). A number of previous studies have investigated the impact of these abiotic environmental parameters on the *Artemia* hatching performance (Ahmed et al., 1997; Bahr et al., 2021; Hasan and Rabbane, 2018; Kumar and Babu, 2015; Sharahi and Zarei, 2016; Wasonga and Olendi, 2018) which is evaluated through the measurement of the hatching rate at the endpoint of the experiment. For instance, Kumar et al. observed that optimal hatching performance of *Artemia* occurred at 29°C and 29 parts per thousand (ppt) salinity (Kumar and Babu, 2015) while Sharahi et al. found the optimal conditions to be 27-28°C and 35 ppt, respectively (Sharahi and Zarei, 2016). Hasan et al. showed that the maximum proportion of *Artemia* hatched when the salinity and temperature were 30 ppt and 24°C, respectively, and that the hatching rate decreased as the temperature rose from 24°C to 32°C while the salinity ranged between 20-40 ppt (Hasan and Rabbane, 2018). Ahmed et al. reported that the optimal salinity for hatchability is 20 ppt (Ahmed et al., 1997). In another study done by Bahr et al. on the influence of salinity on hatching, the optimal hatchability was found at salinities of 60 ppt and temperatures around 30°C (Bahr et al., 2021). The variation in optimal parameters could be due to differences in test settings (e.g., light exposure (Arun et al., 2017; Asil et al., 2023; Sharahi and Zarei, 2016)) or variance in environmental conditions (container size, temperature/salinity gradients in bulk media).

This work aims to unify environmental conditions and extend the understanding of the effects of abiotic parameters beyond the endpoint metric of hatching rate to an in-depth, mechanistic analysis of their impacts on the different stages of the hatching process. To do so, we have integrated an optical oxygen sensor in a microfluidic “aquarium” to monitor rate of oxygen consumption, (also referred to as routine metabolic rate (Killen et al., 2021)) in real-time throughout the entire hatching process. Previously oxygen sensors have been used to monitor respiration in different marine species including fish (Souders et al., 2018; Stokes and Somero, 1999; Warkentin et al., 2007). The relatively small size of the microfluidic platform decreases the signal-to-noise ratio (Curto et al., 2012; Lasave et al., 2015; Qiu and Nagl, 2021) of the sensor and also enables the correlation of sensor data with in situ microscope imaging data. The platform can also more readily maintain uniform environmental conditions (Azuaje-Hualde et al., 2017; Hung et al., 2005; Li et al., 2017) using off-chip programmable controllers rather than bulk systems (such as beakers, fish tanks) used in contemporary studies. Recently, microfluidic and millifluidic lab-on-a-chip platforms have been used for pharmaceutical applications including drug testing on cells (Asif et al., 2020), biological animal studies such as c. elegans (Gokce et al., 2017; Krenger et al., 2020; Mondal et al., 2016; Rahman et al., 2020; Rohde et al., 2007), and zebrafish (Akagi et al., 2013, 2012; Yang et al., 2016). Moreover, The accuracy of the end point analysis of the hatching rate is also enhanced through the automated transfer -minimizing contamination and human error (Dey et al., 2019; Grover et al., 2006; Li et al., 2019; Zhang et al., 2021) -of the entire tested sample to a counting chip with sieve-like structures that can be imaged in its entirety and processed with image analysis rather than relying on a batch sample and manual counting methods.

## 2. Experimental

### 2.1 Artemia cyst

*Artemia* cysts were purchased from Brine Shrimp Direct and stored at 4ºC as recommended by the vendor until one day prior to the hatching studies, at which point they were transferred to room temperature to allow for gradual temperature adaptation. Fig. 1(A) shows the scanning electron microscopy (SEM) (JEOL FS-100) images of the as-received *Artemia* cysts. The cysts are cup-shaped in structure, and the diameter of the circular structure of the cup ranges between 180-260 μm.

**Fig.1.**
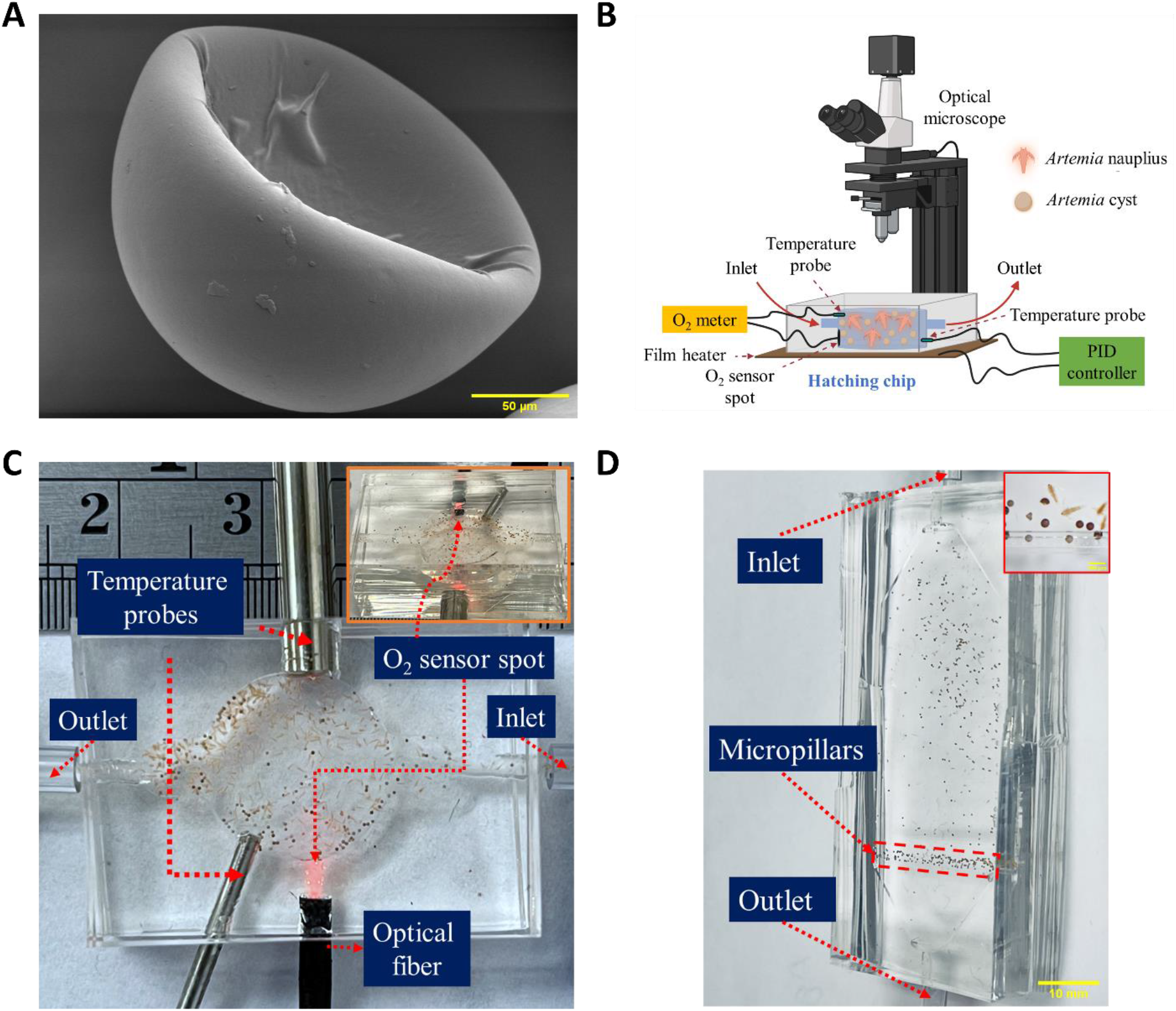
Experimental setup. (A) Scanning electron microscopy (SEM) shows the cup-like structure of the as-received *Artemia* cysts at 450X magnification. (B) Schematic overview of the microfluidic platform to study the effect of abiotic environmental parameters on the hatching performance of *Artemia*. (C) Hatching chip with integrated O_2_ sensor and temperature probes connected (scale bar= 5 mm) (Position of O_2_ sensor spot is shown in inset) (D) Counting chip with sieving-like structures made of PDMS micropillars at the outlet (scale bar= 10 mm) [Cysts and nauplii entrapped at the sieving structures are shown in inset (scale bar= 500 μm)].

### 2.2 Hatching of *Artemia* cysts and automatic hatching performance calculation

The hatching studies of the *Artemia* cysts were performed on a microfluidic platform (Fig. 1B). The platform consisted of a hatching chip, which has a cylindrical chamber (diameter of 13 mm, height of 3.55 mm) connected to an inlet and outlet (height of 1.7mm, width of 4mm) (Supplementary Fig.1A). The total volume of the hatching chip is ∼563 μL. Rounded corners (fillets) were incorporated at the inlet and outlet to ensure cysts were not trapped in the chamber corners during insertion at the starting of the hatching and the transfer of the cysts and nauplii at the end of the experiment. Dead volume was minimized through simulating the fluid flow in ANSYS FLUENT (Supplementary Fig.1B). Fig. 1C shows the hatching chamber used for this study. The chip was composed of polydimethylsiloxane (PDMS) and fabricated by casting PDMS on a mold (Supplementary Fig.1C) prepared using a 3D printer (MAX X-43, Asiga) and cured overnight at 60°C in a gravity convection oven (model: 10GC, Quincy Lab). The hatching chip has two layers. The top layer has the chamber and channel structures, and the bottom layer is a blank PDMS slab. Both top and bottom layers of the hatching chip were bonded using corona treatment (BD-20AC, ETP), and subsequent heating at 60°C for overnight. Before bonding the layers, several holes were made in the top layer using biopsy punches for the inlet, outlet, and sensors. An O_2_ sensor spot (OXSP5, Pyroscience) was bonded to the PDMS top layer inside the chamber using silicone glue (Spglue, Pyroscience). An optical fiber was connected exactly behind the O_2_ sensor spot through a hole in the PDMS chip. The optical fiber was connected to an external optical oxygen meter (FireSting®-O_2_, Pyroscience). The oxygen meter was composed of a light-emitting diode (LED) and a photodiode that both stimulate and detect the oxygen-sensitive spot oxygen-dependent luminescence emission and measures the dissolved oxygen concentration of the water in the chamber every second during hatching. A temperature probe (Pt100, Pyroscience) was also connected to the chip to detect the temperature of the water inside the chip and thus compensate the dissolved oxygen concentration readings for any fluctuation in temperature. To minimize variation, the chip was placed on top of a film heater (PI film heater-24 V, Icstation) connected to a proportional–integral–derivative (PID) controller (6-30V DC Electronic Thermostat Controller, DROK) which controlled the temperature of the water inside the hatching chip according to the set temperature (20, 35 and 30ºC). A variable voltage DC power adapter (ALP002, KEJIESHENG) was used to power the PID controller, and a thermocouple probe (TC 10K, QINGDAO) was connected to it for measuring the water temperature inside the chip. The PID controller adjusts the input power according to the prob measurement to maintain a maximum variation of 0.1°C from the set temperature. The hatching chip with the embedded heater was placed under a digital stereo optical microscope (SE400-Z, Amscope) equipped with a digital camera (MD500, Amscope) to capture a photomicrograph of the chip every five minutes duringhatching. During the entirety of the hatching process, a light-emitting diode (LED) light (1W, Amscope) was kept on continuously due to its effect on hatching (Asil et al., 2023; Kumar and Babu, 2015).

Prior to the start of the experiment, the water for hatching was prepared by adding varying amounts of commercially available sea salt (Fluval Sea, pH: 8.1-8.2) to deionized water (DIW) (prepared using Barnstead Smart2Pure Water Purification System, Thermo Scientific) to produce water with 0, 25, 50, and 75 parts per thousand (ppt) salinities and thoroughly mixed in a test tube using a vortex mixer (Vortex genie 2, Scientific Industries). The appropriate weight of *Artemia* cysts was measured using a digital precision balance (Bonvoisin) and mixed carefully with the saltwater solution so that the cyst concentration was 5 g/L. The cyst solution was immediately inserted inside the hatching chip until the chamber was filled. The inlet and outlet of the chip was then closed using flexible tubes and binder clips to prevent air infiltration inside the chip. Hatching experiments were conducted for 24 hours from the time cysts were immersed in the saline solutions.

The depth of the hatching chip (3.55 mm) was designed to ensure the suspending fluid had sufficient dissolved oxygen for hatching and space for the nauplii to swim. This depth allowed multiple nauplii and cysts in the same vertical plane, which precluded accurate counting of nauplii and cysts. Hence, a counting chip (Fig. 1D) was designed with a shallow depth (800 μm) chamber such that nauplii and cysts were restricted to a single monolayer. The chip had an inlet and an outlet. The outlet contained PDMS micropillars which acted as sieving-like structures (Fig. 1D inset) and allowed only the water to flow through the channel, preventing the nauplii and cysts in the chamber from escaping. The chip had two layers: the top layer had a chamber, micropillar structures, inlet, and outlet and was constructed of PDMS cast on a 3D printed mold (Supplementary Fig.2A). The bottom layer was a blank PDMS slab. Both layers were bonded using corona treatment and subsequent heating, as described above. At the inlet and outlet of the counting chip, the height is higher than that of the chamber (1.7 mm vs. 800 μm) and designed with a fillet radius of 900 μm to allow smooth flow of *Artemia* cyst and nauplii through the inlet and outlet. Supplementary Fig.2B displays the results of an ANSYS Fluent simulation of the fluid flow through the chip, demonstrating the absence of any dead volume. After 24 hours, the hatched nauplii and hatched/unhatched cysts in the hatching chip were transferred to the counting chip automatically in a 30% methanol solution at a 100 μL/min flow rate by syringe pump (Pico plus 11 elite, Harvard Apparatus). The outlet of the hatching chip and the inlet of the counting chip were connected by a PTFE tube, and the inlet of the hatching chip is connected to the syringe mounted in syringe pump via a flexible Tygon tube. *Artemia* nauplii swim rapidly after hatching and the 30% methanol solution in DIW was used to euthanize the nauplii to obtain clear images in the counting chip. A digital camera (D3100, Nikon) was used to capture images of the nauplii, and cysts present in the hatching chip. Images were processed in ImageJ where the number of cysts and nauplii were counted based on their circularity (Fig. 1E). If any object had a circularity greater than or equal to 0.9, it was considered to be a cyst (hatched or unhatched), and any object in the image with lower circularity (less than 0.9), was considered to be nauplii. The hatching rate was calculated using the following formula-

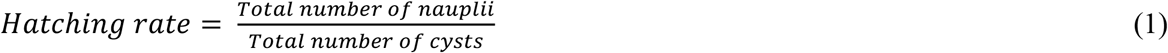

### 2.3 Statistical analysis

OriginPro (ver. 2022b, OriginLab) was used for all statistical analysis. Data are represented as the mean ± standard deviation. The significance of data within and between different groups was measured using two-way analysis of variance (ANOVA) with Bonferroni post-hoc test when all categories of data were available [hydration and differentiation stage (discussed later)], if not, separate one-way ANOVA with Bonferroni post-hoc test was performed [emergence and hatching stage (discussed later)]. Analysis of the correlation between different results was performed using the Pearson correlation coefficient. Data were considered significant, if *p*<0.05.

## 3. Results and discussion

### 3.1 On-chip detection of oxygen consumption and morphological changes in cysts during the hatching process

The hatching process of *Artemia* can be divided into four stages: i) hydration, ii) differentiation, iii) emergence, and iv) hatching (Emerson, 1967), where each stage is characterized by a different energy metabolism (Drinkwater and Crowe, 1991; Emerson, 1963). Fig. 2A shows the photomicrographs of different stages of hatching. And in Fig. 2B, we demonstrate that each of these stages and the transition between them can be identified through changes in depletion in dissolved oxygen concentration (DDOC) of the water in the hatching chip and the measured rate of oxygen consumption (ROC) where ROC is defined as the change in total DDOC divided by the duration of the respective stage. It is important to note that although PDMS is permeable to oxygen (Rao et al., 2007; Shiku et al., 2006), comparison of the measured data (solid lines, Fig 2B) to the control measurement of the DDOC level without cysts indicates (dotted lines, Fig 2B) that the rate of consumption at all stages is significantly larger than permeation thereby supporting the use of the present setup to qualitatively compare the DDOC under different environmental conditions.

**Fig. 2.**
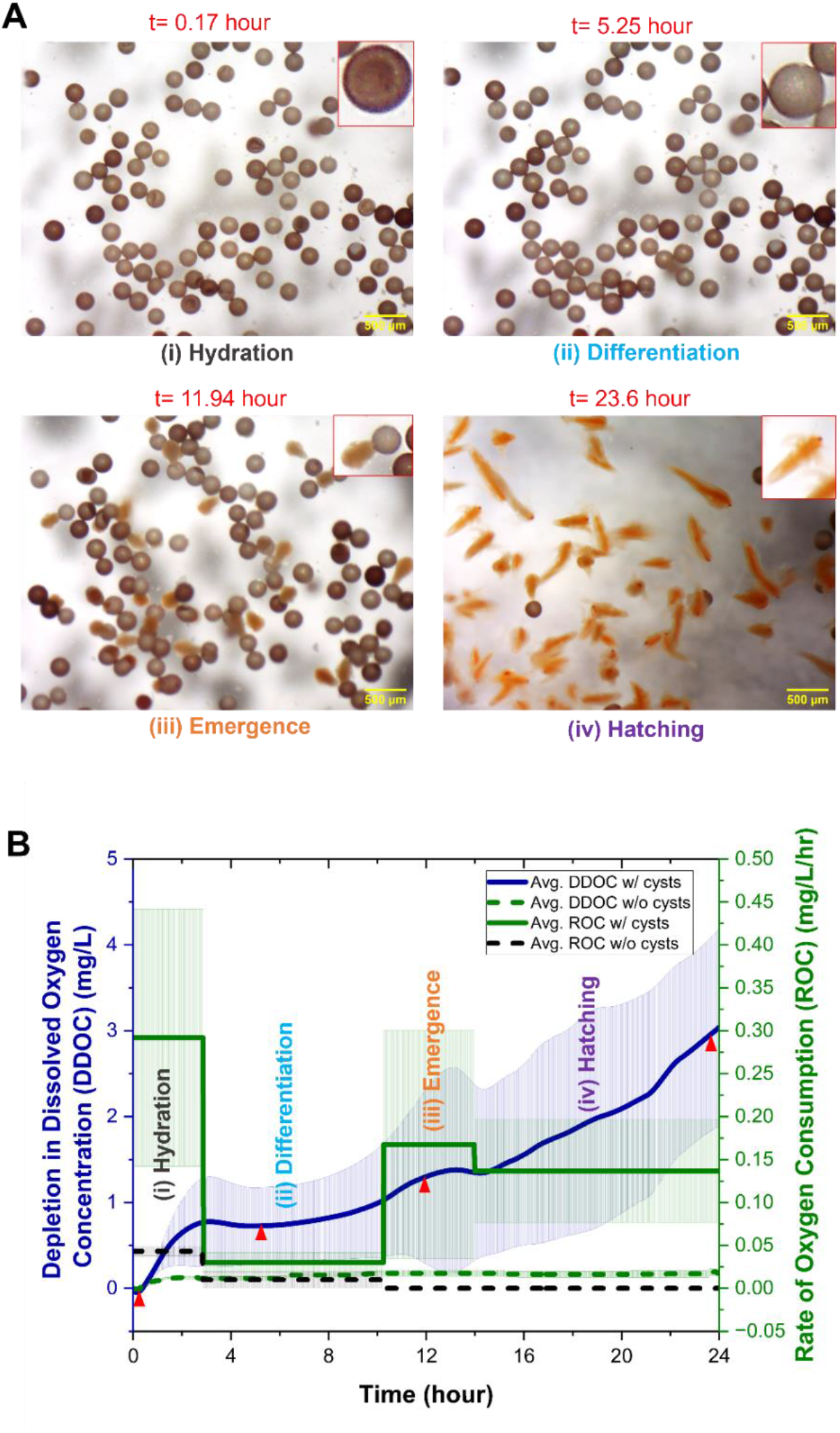
Photomicrographs and O_2_ sensor readings at different stages of hatching. (A) Photomicrograph at different stages of hatching obtained from the optical stereomicroscope (the inset shows the morphology of the cysts/nauplii at different stages). (B) On-chip detection of the change in oxygen consumption behavior of the *Artemia* cysts at different stages of hatching through the depletion in dissolved oxygen concentration (DDOC) and rate of oxygen consumption (ROC) at 25 ppt water salinity, and temperature of 25°C (the bold lines in the graph show the average and the shaded area shows the standard deviation). Red triangles in Fig. 2(B) represent the DDOC at the time points corresponding to the photomicrographs presented in Fig 2(A).

Immersion of the cysts in water starts the hydration stage, in which the cysts inflate and develop into a spherical (rather than cup like) shape [Fig. 2A(i)]. As the cysts imbibe water, energy metabolism, RNA, and protein synthesis begin within a short period (Clegg and Trotman, 2002; Emerson, 1963). As energy metabolism is reactivated, respiratory activity increases dramatically, as evidenced by the increased ROC [Fig. 2B(i)]. At the end of the hydration stage, the majority of the cysts are spherical in shape and differentiation commences. At this stage, there is no cell division and no increase in DNA (Clegg and Trotman, 2002; Emerson, 1963) resulting in an almost constant oxygen requirement and thus lower ROC [Fig. 2B(ii)]. In the third stage (emergence), the cyst shell starts to fracture due to the increased turgor pressure within resulting from an in creased amount of glycerol, and the embryo starts to emerge from the broken shell within a hatching membrane [Fig. 2A(iii)] (Drinkwater and Crowe, 1991). The emergence stage is highly active, resulting in an increase in DDOC and ROC [Fig. 2B(iii)]. Finally, at the last stage (hatching), the *Artemia* embryos leave the cyst shell and hatching membrane and begin to swim [Fig. 2A(iv)]. At this stage, oxygen demand lowered again, as detected by a comparative decrease in ROC [Fig. 2B(iv)].

### 3.2 Effect of water salinity and temperature on the duration of different stages of hatching

In Fig. 3, the variation in the duration of each stage of hatching under varying temperatures and salinities is obtained through measured changes in the ROC and verified through comparison with photomicrographs. Overall, we note that at low temperatures and low salinities (with the exception of 25 ppt at 20°C and 0 ppt at 30°C), the 24-hour time period was insufficient to complete the hatching cycle due to the increased duration of each of the other three stages. In line with this, the duration of the hatching stage (defined as the time remaining at the end of the emergence stage) was found to increase considerably when salinity changed from 0 to 75 ppt salinity and to significantly decrease at 20°C.

**Fig. 3.**
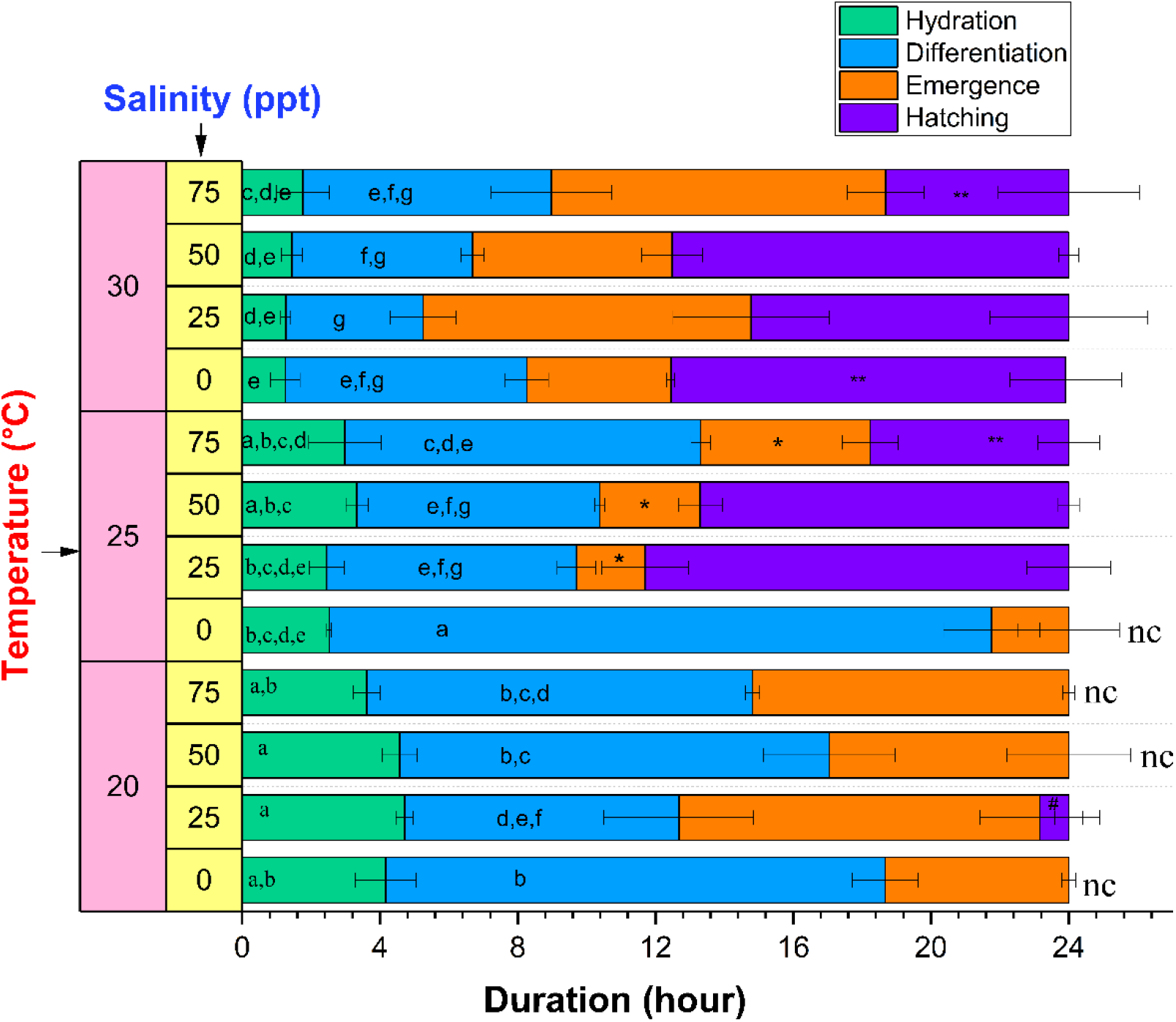
Duration of different stages of hatching at different water salinities and temperature. Values are expressed as mean ± standard deviation (n=3). In each category (hydration, differentiation, emergence, or hatching stage duration), means that do not share the same letter are significantly different from each other (two-way ANOVA with Bonferroni post-hoc test, *p*<0.05). The hatching stages which were not complete within 24 hours were not statistically analyzed and marked with “nc”. *, and # denote significantly different emergence and hatching stage duration, respectively, at a temperature as compared to the other tested temperatures (one-way ANOVA with Bonferroni post-hoc test, *p*<0.05). ** denotes significantly different hatching stage duration between two different tested salinities (one-way ANOVA with Bonferroni post-hoc test, *p*<0.05).

Recognizing that during hydration (green bars), the dry *Artemia* cysts absorb water by osmosis (CLEGG, 1964), the effect of temperature and salinity can be modelled by Van’t Hoff’s law (Shin and Kim, 2018),

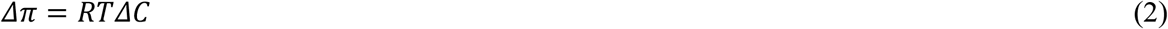

where Δπ is the difference in osmotic pressure, R is the universal gas constant, T is the temperature, and ΔC is the difference in solute concentrations. As the temperature (T) increases, so does the rate of osmosis, corresponding to a decrease in the hydration duration as illustrated in Figure 4.

**Fig. 4.**
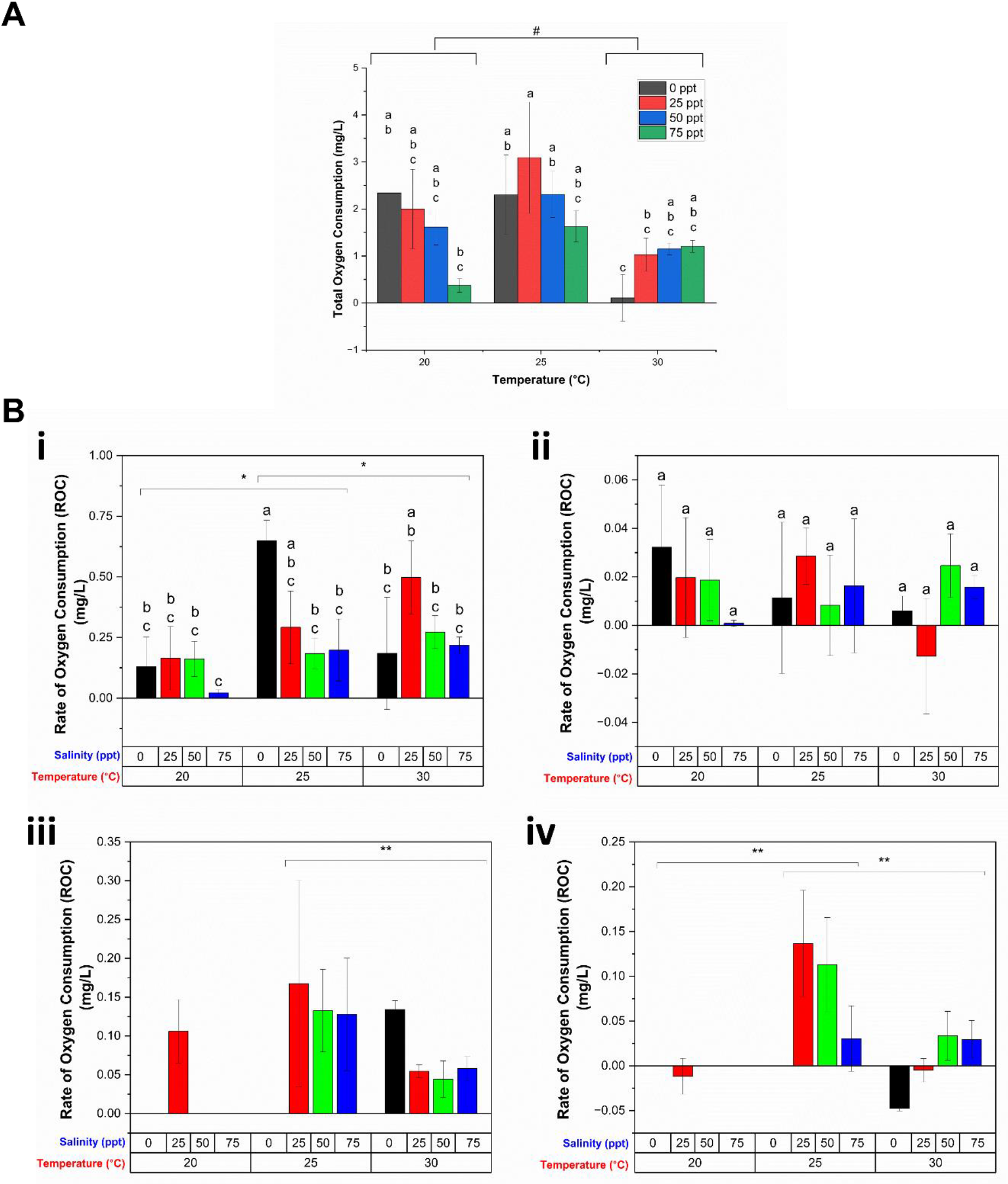
Oxygen consumption of the *Artemia* cysts under different temperatures and salinities. (A) Total oxygen consumption, or total DDOC throughout the entire hatching process under different temperature and salinities. (B) Rate of oxygen consumption (ROC) at different stages of hatching: i) hydration, ii) differentiation, iii) emergence, and iv) hatching stage. Values are expressed as mean ± standard deviation (n=3). In each category of data (total oxygen consumption, or ROC in hydration and differentiation stages), means that do not share letters above the columns are substantially different from each other (two-way ANOVA with Bonferroni post-hoc test, *p*<0.05). At different tested temperature, statistically significant ROC was indicated by * (two-way ANOVA with Bonferroni post-hoc test, *p*<0.05) and ** (one-way ANOVA with Bonferroni post-hoc test, *p*<0.05) respectively, whereas # indicates statistically significant total oxygen consumption (two-way ANOVA with Bonferroni post-hoc test, *p*<0.05).

Supplementary Fig. 3A shows the significant negative correlation between the hydration duration and temperature determined by the Pearson correlation coefficient (r=-0.9) and a linear fit between these parameters (R^2^=0.99) (Supplementary Fig. 3B). At the same time, the increase in water salinity at any temperature can decrease the difference in internal and external solute concentrations (ΔC) of cysts, resulting in a decrease in osmosis and oxygen consumption with an increase in the hydration duration. Interestingly, the present results indicate that changes in the salinity alone is not sufficient to significantly increase duration, but this increase was more pronounced as the temperature decreases. On the other hand, the duration of differentiation (blue bars, Figure 4) decreases with both increased salinity and temperature, except at very high salinities. To understand this, we note that through the differentiation stage, the cyst reaches the emergence stage, when the increasing turgor pressure induced by the synthesis of glycerol from trehalose causes the shell to begin to fracture. A rise in temperature enhances glycerol synthesis (CLEGG, 1964), which in turn shortens the period of differentiation. Increasing salinity also increases the glycerol level (Drinkwater and Crowe, 1991), and may decrease the differentiation period. However, as water salinity rises, the external environment becomes hypertonic, and there is a chance of cellular water loss due to the exosmosis process, which could lengthen the period of differentiation at higher salinity.

Interestingly, the emergence stage duration (orange bars) suggests an optimum temperature of 25°C with a significant decrease in duration, compared to that at 20 and 30°C. Although temperature helps to increase the osmotic pressure and thus reduces the hydration and differentiation stage length, once the emergence stage starts, and the embryo emerges within the hatching membrane, it appears to have less tolerance for higher temperatures. Hence, an intermediate temperature of 25°C was more favorable for the emergence stage and required less time for completion.

### 3.3 Effect of water salinity and temperature on oxygen consumption during hatching

Further understanding of the hatching process is obtained by directly examining the depletion in dissolved oxygen concentration (DDOC) of water in the hatching chamber for varying temperatures and salinities. Supplementary Fig. 4 shows the average DDOC curves of triplicate experiments performed at each temperature and salinity condition. Overall, the results indicate a positive correlation between hatching duration and DDOC for salinities greater than zero. The total DDOC, or total oxygen consumption, for the 24-hour hatching period is shown in Fig. 4A, indicating the presence of an optimum temperature (25°C), because total oxygen consumption is higher at this temperature regardless of salinity. For all temperatures, the total oxygen consumption was reduced between 25 to 75 ppt. The interactive effect of temperature and salinity on the total oxygen consumption was also statistically significant.

A more detailed understanding of the effects of temperature and salinity on the hatching process is obtained through consideration of oxygen consumption at each of the four stages of hatching. Since the duration of different stages of hatching varied with temperature and salinity, the results are normalized by considering the rate of oxygen concentration (ROC) at each stage (Fig. 4B), which can be also considered as standard metabolic rate as mentioned in the literature (Killen et al., 2021). During hydration, ROC increased significantly with temperature irrespective of salinity [Fig. 4B(i)]. ROC also significantly decreased with increased salinity between 25-75 ppt which reflects lower activity in accordance with the longer duration (Figure 3). On the other hand, although both salinity and temperature decreased duration of differentiation, the metabolic rate was not significantly affected. (Fig. 4B(ii)). During the emergence stage, ROC was found to be significantly affected by temperature, but not salinity at *p*= 0.05 level (one-way ANOVA), with ROC being significantly decreased when temperature was increased from 25 to 30°C (Bonferroni, *p* < 0.05, Fig. 4B(iii)). We note that the statistical significance of the data was assessed using one-way ANOVA with Bonferroni post-hoc test due to incomplete emergence in some experimental conditions (Figure 3). The decreased oxygen consumption with increased temperature may indicate physiological stress. Finally, in the hatching stage, ROC is maximum at 25°C, and decreases with increasing salinity.

### 3.4 Effect of water salinity and temperature on hatching rate

Figure 5 illustrates the hatching rate of the *Artemia* cysts hatched under different temperatures and water salinities in our platform. In accordance with previous work, the results indicate the existence of optimal conditions (Hasan and Rabbane, 2018; Sharahi and Zarei, 2016). Specifically, the hatching rate increased dramatically when the temperature rose from 20 to 25°C, but there was no significant change as the temperature rose further. Similarly, the hatching rate increased when salinity was increased from 0 to 25 ppt, but subsequently reduced with increased salinity. A maximum hatching rate was observed at 25°C temperature and 25 ppt water salinity (76.85±14.22%) in accordance with the measured maximum ROC (Figure 4iv). Moreover, the measured ROC indicated that the very low hatching rates at zero salinity and 20°C do not necessarily reflect hostile conditions (since we do observe activity) but that the 24-hour duration is insufficient.

**Fig. 5.**
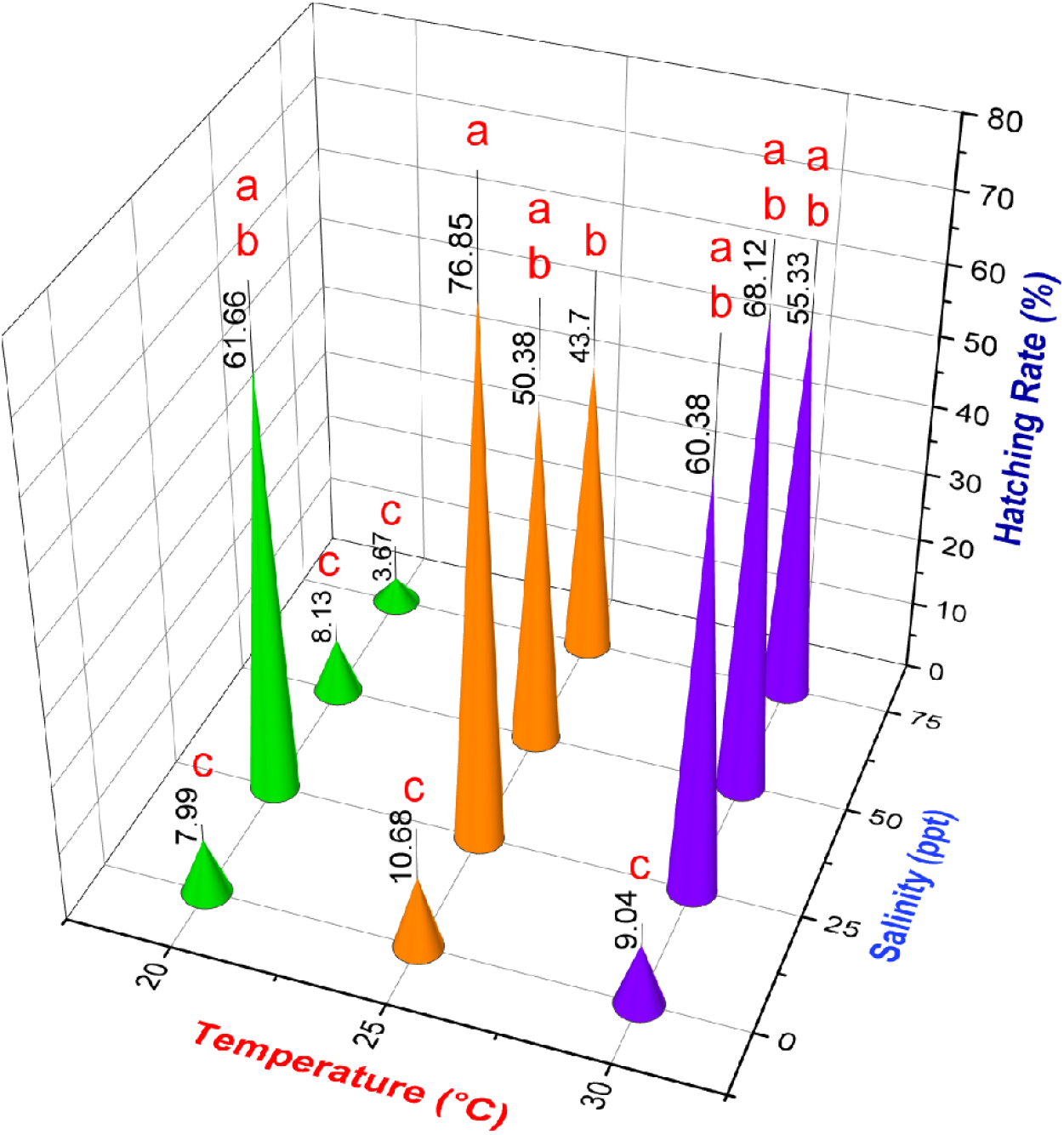
On-chip hatching rate calculation of *Artemia* at different water salinities and temperatures. Values are expressed as mean ± standard deviation (n=3). Means that do not share the same letter above the columns are substantially different from each other (Two-way ANOVA with Bonferroni, *p*<0.05).

### 3.5 Correlation of oxygen availability, respiration behavior, duration of different hatching stages and hatching rate

The present approach, in which the entire hatching process is monitored in real time, uniquely advances the understanding of the relationships between the abiotic parameters, stage performance (duration, ROC) and hatching rate. To quantify these parameters, we implement a Pearson correlation (Fig. 6). We first note that although ROC was significantly affected by temperature, salinity, or both, only ROC at the hatching stage had a statistically significant association (positive correlation) with the hatching rate (Fig. 6A). On the other hand, only differentiation duration is significantly correlated with hatching rate (negative correlation) (Fig. 6B).

**Fig. 6.**
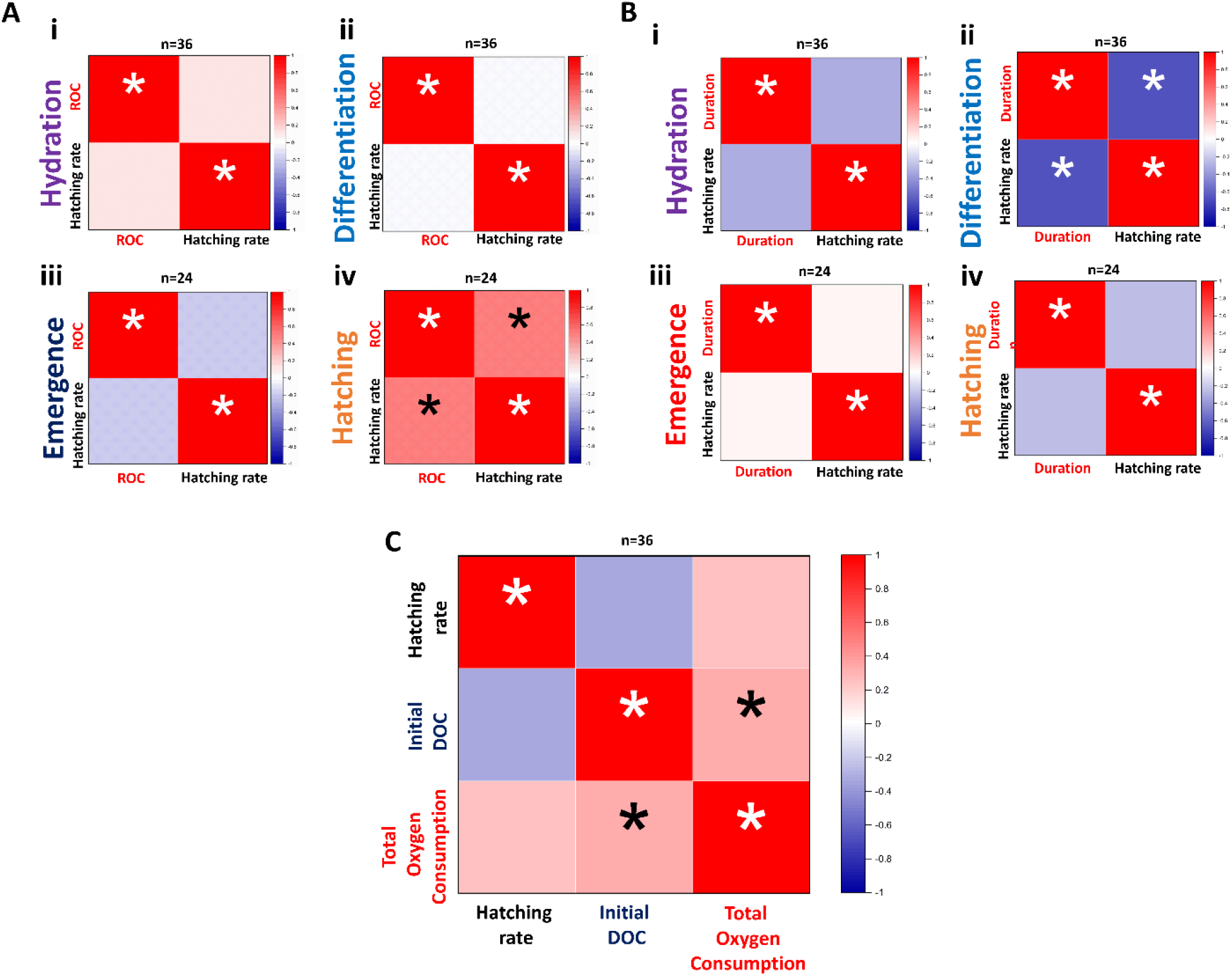
Pearson correlation factor analyzed between- (A) rate of oxygen consumption (ROC) at different stages and hatching rate, (B) duration of different stages and the hatching rate, (C) hatching rate, initial dissolved oxygen concentration (initial DOC), and total oxygen consumption [* represents relation is statistically significant (*p*<0.05)].

Negative correlations between hatching rate and hydration, and differentiation stage duration and a positive correlation between hatching rate and hatching stage duration were observed (Fig. 6B). The hatching rate was maximum when the temperature and salinity was 25ºC and 25 ppt, respectively. At lower or higher temperature or salinity among the tested conditions, the hydration and differentiation stage took longer, and hatching stage was shorter. Similarly, oxygen consumption correlates positively with hatching rate at the hydration and hatching stage, but negatively with hatching rate at the differentiation stage. At 25ºC temperature and 25 ppt salinity, the oxygen consumption was comparatively high at the hydration stage, lower during the differentiation stage, and very high at the hatching stage. This pattern was reversed at very low or very high tested temperatures or salinities. This suggests as temperature and salinity significantly affect the whole hatching process, higher hatchability is obtained at that favorable temperature and salinity condition, when the majority of the cysts become hydrated and regain a higher metabolic rate in a shorter period of time during the hydration stage, expend less energy during the differentiation stage and complete it more quickly. At this optimal condition, cysts will spend longer time in the hatching stage, resulting in a greater number of cysts hatching and using more oxygen.

Finally, since the concentrations of dissolved oxygen molecules in saltwater varies with temperature and salinity, it is vital to establish if oxygen availability may hinder hatchability (Moens et al., 2018). To this end, the Pearson correlation coefficient was used to calculate correlations between the *Artemia* hatching rate, initial dissolved oxygen concentration (initial availability of oxygen), and total oxygen consumption (Fig. 6C). Although the initial dissolved oxygen concentration (initial DOC) had a significant positive correlation with the total oxygen consumption, it had a negative nonsignificant correlation with the hatching rate. This means within our tested conditions, *Artemia* cysts could consume more oxygen based on the higher availability of oxygen (at low temperature and salinity); however, this is not indicative of a greater number of *Artemia* hatching. Depending on the availability of oxygen, *Artemia* cysts might likely acclimatize; nonetheless, the interaction between temperature and salinity governs hatchability.

## 4. Conclusion

In this work, we have demonstrated that real time measurement of metabolic rates may offer in-depth, mechanistic understanding of ***how*** abiotic parameters affect marine organisms. Focusing on temperature and salinity (two significant environmental parameters that are also susceptible to climate change) and a major zooplankton, *Artemia* as a model organism, we measure the respiration behavior, progression of hatching as well as the resulting number of hatched *Artemia* under distinct temperature and salinity levels throughout a predetermined time period.

In accordance with previous studies, extreme salinities (low or high) and low temperatures are observed to inhibit *Artemia* hatching verifying the existence of optimal hatching conditions (Ahmed et al., 1997; Bahr et al., 2021; Hasan and Rabbane, 2018; Kumar and Babu, 2015; Sharahi and Zarei, 2016; Wasonga and Olendi, 2018). However, while these previous studies primarily focus on endpoint measurements of the hatching rate, here we have established relationships between the hatching rate and the progression of hatching and respiration behavior throughout the process. For example, we demonstrate that a key factor affecting the hatchability of *Artemia* in a 24h time period is the duration of the differentiation stage, which is found to be inversely proportional to the hatching rate. Overall, the duration of differentiation (blue bars, Figure 4) decreases with both increased salinity and temperature, except at very high salinities. Based on an understanding of the differentiation stage, it is possible that this relationship is connected to increased glycerol production that accelerates shell fracture characterizing the onset of emergence. Although validation of this hypothesis is left to future work, we illustrate that the current experiments can highlight key areas of focus towards understanding the complex relationship between environment and marine health. A crucial link between temperature, salinity and the reactivation of latent cyst metabolism was also clearly demonstrated by comparing hydration progression with Van’t Hoff’s law.

The present experimental approach was enabled through development of a novel microfluidic platform with integrated oxygen sensor, in which physical changes in the *Artemia* cysts could also be recorded using an optical microscope. Precise environmental control is obtained using a heater with a feedback loop and an automated counting chip is designed to eliminate errors associated with the lack of stringent environmental control and manual calculation of hatchability. As well as offering a more in-depth look at biological processes, widespread use of automated systems with precise environmental control standardizes experiments resulting in more informative cross-comparison in the literature. Thus, the present approach is anticipated to have broader applicability in the research of zooplankton and fish larvae including under varied abiotic/biotic environmental conditions and aquatic contaminants.

## CRediT authorship contribution statement

**Preyojon Dey:** Conceptualization, Methodology, Investigation, Formal analysis, Visualization, Writing - Original Draft **Terence M. Bradley:** Conceptualization, Supervision, Formal analysis, Writing - Review & Editing, Funding acquisition **Alicia Boymelgreen:** Conceptualization, Supervision, Formal analysis, Writing - Review & Editing, Funding acquisition

## Acknowledgments

This work is supported by the National Science Foundation (award number: 2038484, year: 2020). Graphs were plotted using OriginPro 2022b (https://www.originlab.com/). Schematics were created using Biorender (https://biorender.com/).

