## Supplementary information for "The impact of selected abiotic factors on zooplankton hatching process through real-time, in-situ observation"

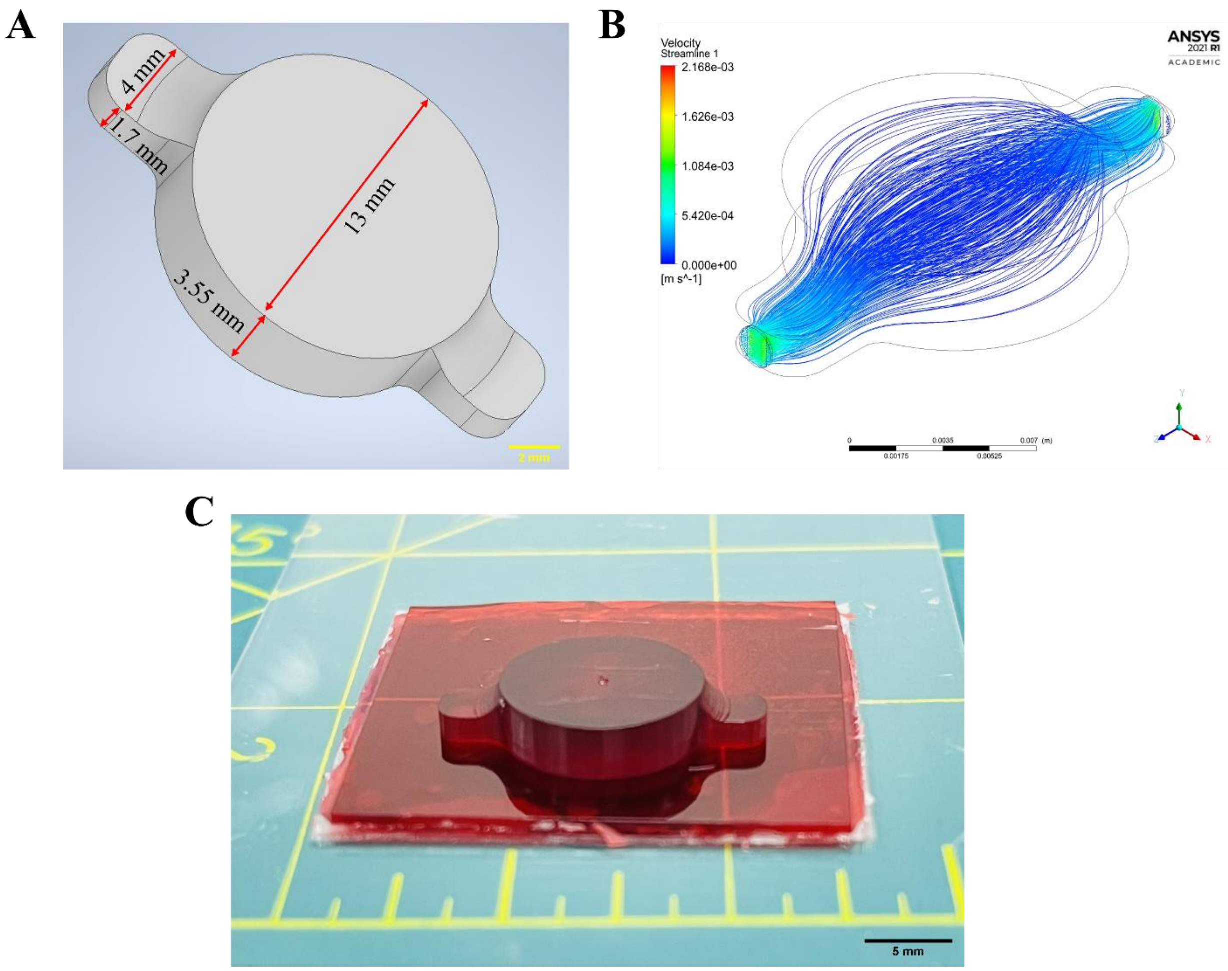


**Supplementary Fig.1** (A) CAD model of the hatching chip. (B) Fluid flow simulation in the hatching chip using ANSYS FLUENT (flow rate=100 μL/min). (C) 3D printed mold for hatching chip.

A

B

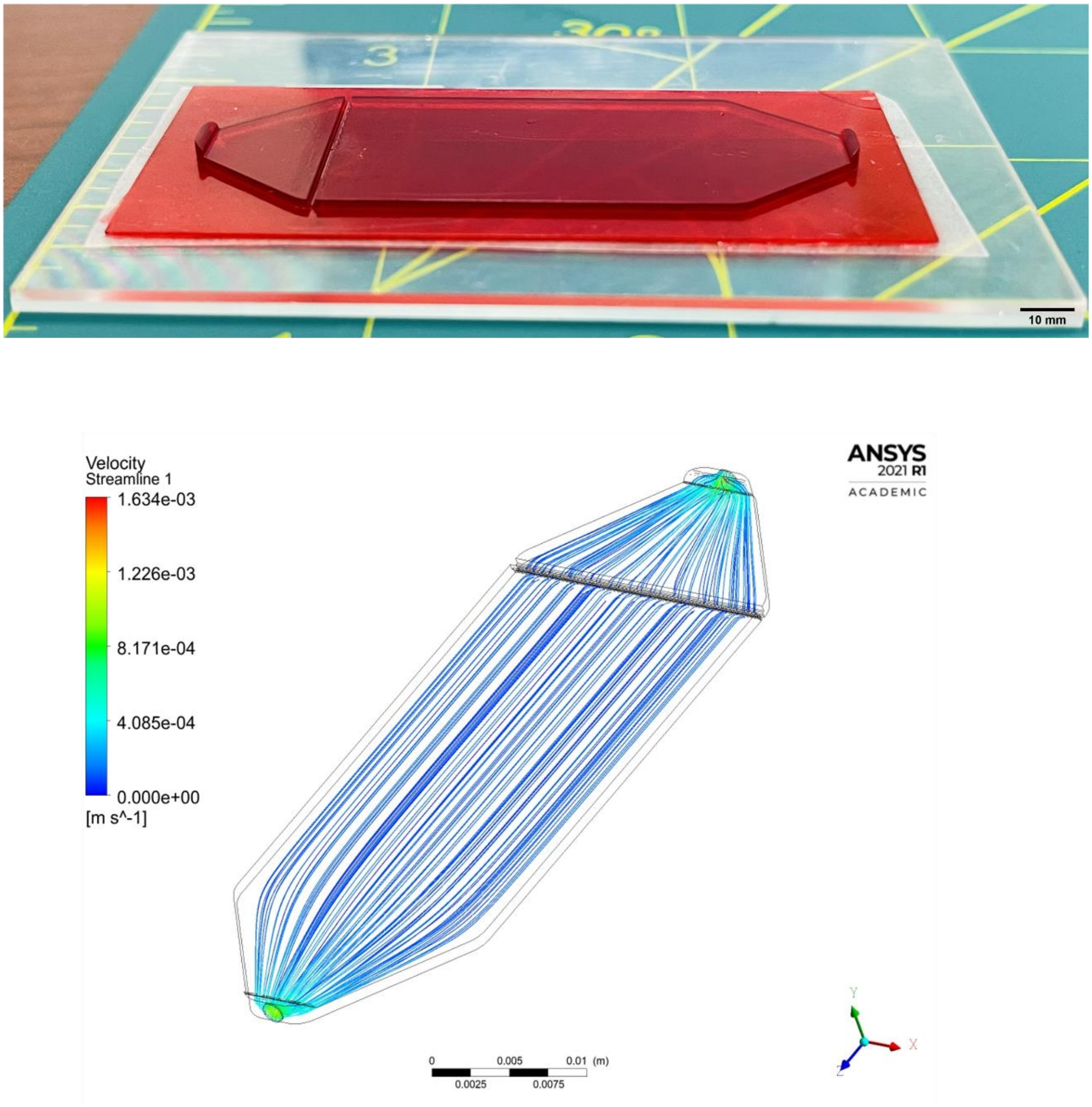


**Supplementary Fig.2** (A) 3D printed mold for counting chip, (B) Fluid flow simulation of the counting chip performed on ANSYS Fluent (flow rate=100 μL/min).

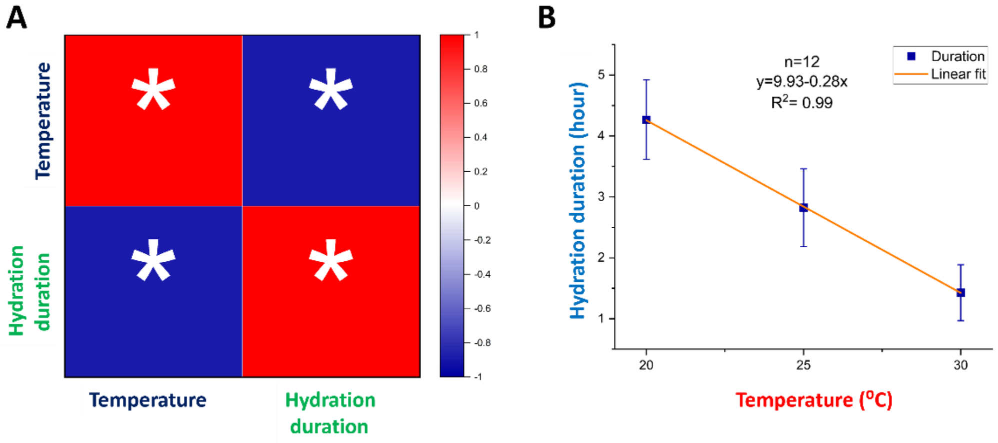


**Supplementary Fig.3** Relation between temperature and hydration duration- A) Pearson correlation coefficient [n=36, * represents relation is statistically significant ($p$<0.05)]. B) Linear fit between hydration duration and temperature irrespective of salinity. Values are expressed as mean ± standard deviation (n=12 at each temperature point).

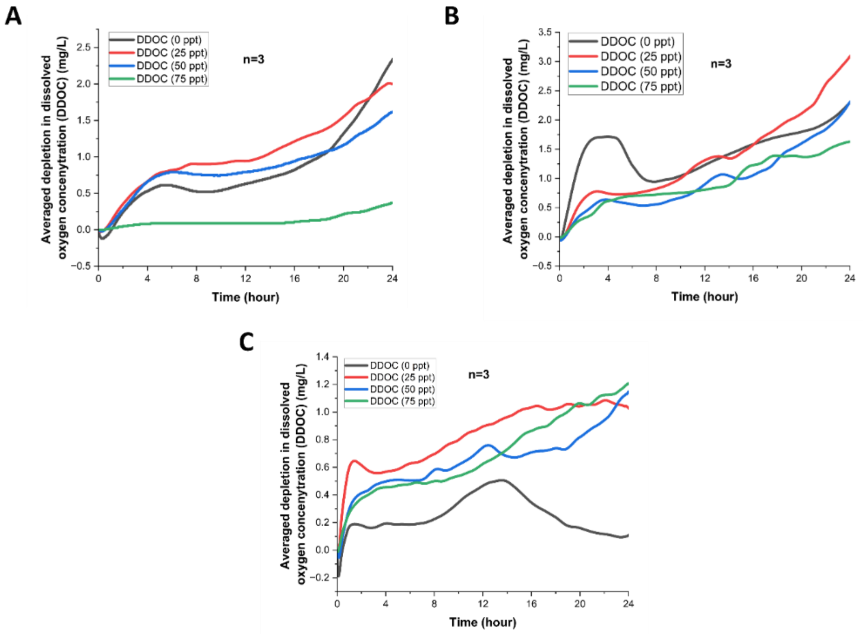


**Supplementary Fig.4** Averaged changes in the depletion in dissolved oxygen concentration (DDOC) due to the oxygen consumption during hatching of *Artemia* under different salinities (0, 25, 50, and 75 ppt) and temperatures of A) $20^0$C, B) $25^0$C, and C) $30^0$C (n=3).
